# PAMPHLET: A Robust Toolkit for Precise PAM Prediction and Unveiling PAM Consistency in Highly Co-occurrence CRISPR-Cas Systems

**DOI:** 10.1101/2024.04.09.587696

**Authors:** Chen Qi, Xuechun Shen, Baitao Li, Chuan Liu, Lei Huang, Hongxia Lan, Donglong Chen, Yuan Jiang, Dan Wang

## Abstract

The CRISPR-Cas technology has sparked a new technological revolution, significantly enhancing our ability to understand and engineer organisms. The nuclease that underpins this technology is evolving from the “One Cas9 for all” model to a diverse CRISPR toolbox. Identifying PAM sequences is a critical bottleneck in developing novel Cas proteins. Given the limitations of experimental methods, bioinformatics approaches are essential for predicting PAM sequences of Cas proteins in advance. To date, there are only a few PAM sequence prediction programs, and their accuracy is relatively low due to the limited number of spacers in CRISPR-Cas systems. To overcome this challenge, we have developed a pipeline named PAMPHLET, which innovatively utilizes homology searches of Cas proteins to identify additional spacers. PAMPHLET was tested on 20 CRISPR-Cas systems with known PAMs, increasing the number of spacers by up to 18-fold compared to the original datasets and successfully predicting 18 PAM sequences for protospacers. For rigorous and high-quality wet-lab validation of the predictions made by PAMPHLET, we employed the published DocMF platform. This platform leverages next-generation sequencing chips to profile protein-DNA interactions and can simultaneously screen both 5’ and 3’ PAMs with high throughput. The PAMPHLET predictions showed high consistency with the DocMF results for four novel Cas proteins. We expect that PAMPHLET will overcome the current limitations in PAM sequence prediction, expedite the discovery of PAM sequences, and help to shorten the development cycle for CRISPR tools. Remarkably, PAMPHLET has revealed an intriguing genomic phenomenon: the C2c9 and C2c10 systems, which lack the canonical adaptation module, possess identical PAM sequences to those found in co-occurring type I systems, suggesting potential shared spacer acquisition mechanisms. This finding highlights the complex evolutionary relationships of CRISPR-Cas systems and propels us toward a deeper understanding of their mechanistic diversity and adaptability.

## 1. Introduction

CRISPR-Cas (Clustered regularly interspaced short palindromic repeats, CRISPR-associate proteins) system, originally discovered as an adaptive immunity mechanism in bacteria and archaea against bacteriophages and plasmids (Barrangou et al., 2007), has been widely utilized as effective and precise gene editing tools because of its RNA-guided DNA cleavage activities (Komor et al., 2017; Pickar-Oliver & Gersbach, 2019). Over the past decade, CRISPR-Cas has revolutionized the field of genetic engineering and has numerous applications in biotechnology, medicine, and agriculture. Scientists have built efficient platforms for creating knockout animal models (Kang et al., 2019; Musunuru et al., 2021), conducting genetic screening (Doench, 2018), and performing multiplexed editing by leveraging the fundamental activity of CRISPR-Cas to generate a targeted genetic disruption in a gene or gene regulatory element (Abdelrahman et al., 2021; Armario Najera et al., 2019). Programmable genome editing technologies have paved the way for gene therapy to treat genetic diseases such as sickle cell disease (Demirci et al., 2021) and *β*-Thalassemia (Frangoul et al., 2021). Additionally, these technologies have been used in plant breeding to rapidly alter plant traits, such as disease resistance or improved yield (Mao et al., 2019).

While this technology has made significant progress in recent years, there are still limitations to current tools that make genome editing applications challenging. Many common CRISPR-Cas effectors are large in size, requiring over 3 kb of DNA sequence to encode the expression construct. This can further reduce transformation efficiencies, which is less desirable, making it more difficult to edit genome in cells. Furthermore, CRISPR-Cas effectors have the potential to exhibit off-target activities or unexplained toxicities. Cas proteins can still recognize and cut the DNA sequence, even if the sgRNA sequence does not perfectly match the DNA sequence, especially at the distal end of the PAM sequence. This tolerance can affect housekeeping genes and cause vital damage. These limitations also present opportunities to advance the field by providing a more comprehensive toolbox for genetic manipulation (J. Y. Wang & Doudna, 2023). Various novel CRISPR-Cas tools like Ca12f1 (Harrington et al., 2018), CasΦ(Cas12j) (Pausch et al., 2020), Cas13X(Cas13e) (Xu et al., 2021), and Cas*π*(Cas12l) (Sun et al., 2023) have been characterized and applied in animal, plant, and human cells to improve accuracy or reduce system size. Notably, some of these emerging CRISPR-Cas tools, such as C2c9 and C2c10, lack the adaptation module (cas1/cas2/cas4) in their original CRISPR-Cas systems, and the mechanism by which they confer immunity is not yet clear.

The identification and characterization of new CRISPR systems have become more straightforward due to the combination of well-established experimental processes and bioinformatics tools. There are three key components that need to be identified in a novel CRISPR-Cas system: the effector protein, crRNA, and PAM. The experimental processes for protein purification, sgRNA design, and PAM screening have been validated and are reliable. Furthermore, bioinformatics tools are used to analyze and compare new CRISPR systems, accelerating the characterization process by elucidating their similarities and differences with known systems. Effector proteins with various functional domains can be identified through methods such as PSI-BLAST and HHpred (Makarova et al., 2020). Additionally, the highly accurate protein structure prediction capabilities of AlphaFold facilitate the functional analysis of effector proteins from novel systems (Jumper et al., 2021). The canonical crRNA is composed of a partial spacer unit flanked by repeats, forming a hairpin structure typical of RNA molecules (Briner et al., 2014). This secondary structure can be characterized with RNA folding tools like Mfold (Zuker, 2003). In some CRISPR-Cas systems, such as type II and certain type V variants, an additional transactivating crRNA (tracrRNA) is necessary for cleavage activity (Deltcheva et al., 2011). New tools have emerged that assist in locating tracrRNA by searching noncoding regions for the anti-repeat sequence of the crRNA (Chyou & Brown, 2019; Mitrofanov et al., 2022). The PAM, which flanks the protospacer either upstream or downstream, is essential for distinguishing self from non-self DNA and preventing autoimmunity. Determining functional PAM sequences is a rate-limiting step in the characterization of novel CRISPR-Cas families, as traditional *in vivo* PAM screening experiments are time-consuming (Tang & Gu, 2020). Despite the availability of high-throughput *in vitro* PAM determination tools like DocMF based on BGISEQ-500 (Li et al., 2020), the effector proteins and crRNA must be prepared in advance. Thus, a bioinformatics-based PAM prediction method is desirable to streamline the PAM screening process and expedite the characterization of new CRISPR-Cas systems.

The immune mechanisms of CRISPR-Cas systems are well-understood, providing a theoretical foundation for our *in silico* predictions (Figure 1). As part of adaptive immunity, the CRISPR-Cas response unfolds in three main stages. Initially, when phages or exogenous plasmids infect bacteria possessing CRISPR-Cas systems, the Cas1 and Cas2 complex incorporates small DNA fragments from the invaders into the CRISPR array, creating a genetic ‘memory’ of the infection in a process known as adaptation. Subsequently, the CRISPR array is transcribed, and the pre-crRNA is processed to yield mature crRNA featuring a single spacer unit flanked by partial repeats. In the final interference stage, in-dividual crRNA assembles with effector proteins to form crRNA-effector complexes, which target and destroy the complementary DNA sequences when the bacteria encounter the foreign DNA again (Mu et al., 2022). For type I, II, and V CRISPR-Cas systems, the PAM sequence is essential for distinguishing endogenous DNA from heterologous DNA during both the adaptation and interference stages (Gleditzsch et al., 2019). The double-stranded DNA cleavage mediated by Cas nucleases depends on accurate PAM recognition (Xue & Greene, 2021). This specificity acts as a safeguard within the CRISPR-Cas adaptive immunity system, ensuring that nuclease activity is directed solely at foreign genetic elements with the corresponding PAM, thus preventing undesired cleavage of the host genome.

**Figure 1:**
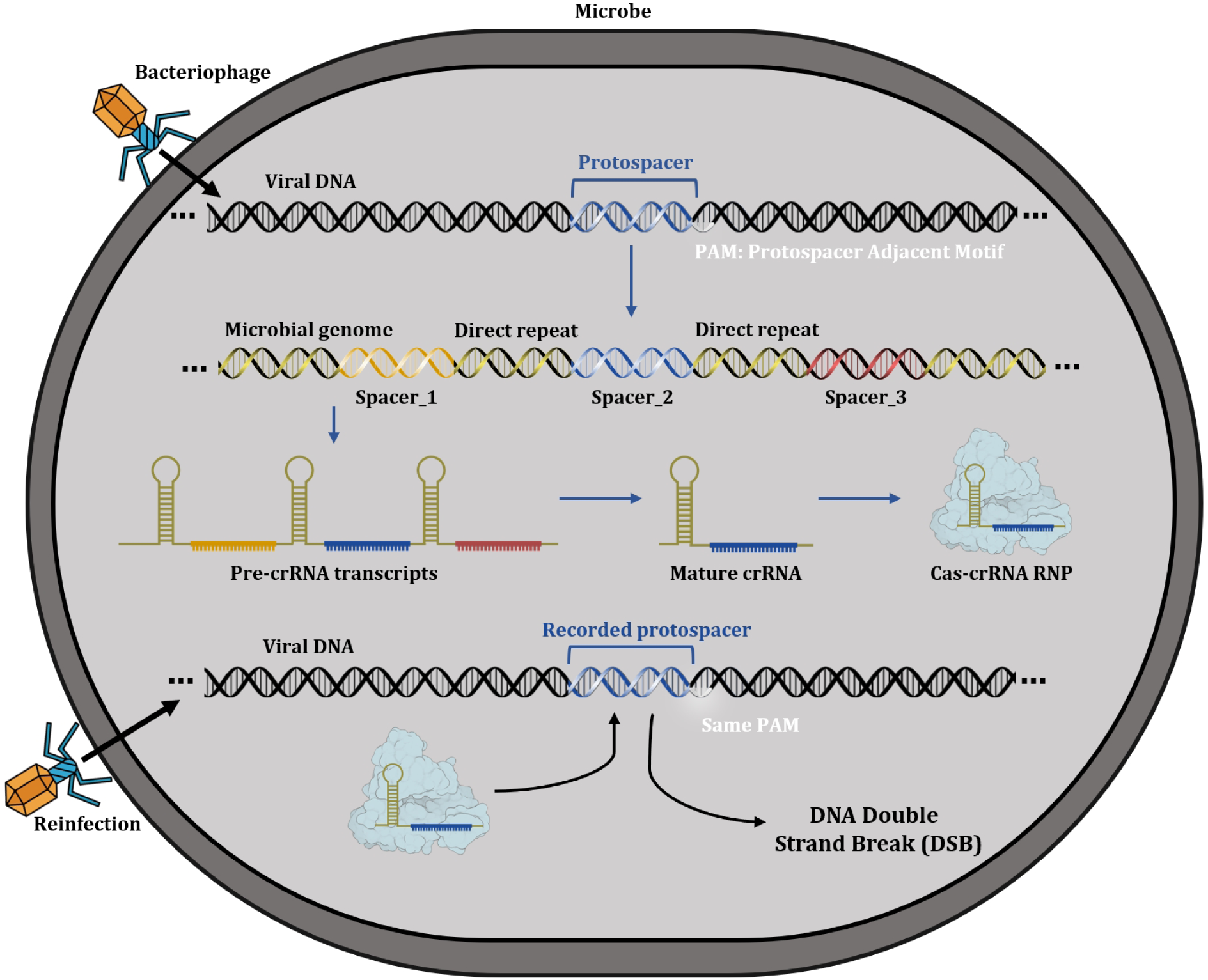
Molecular mechanisms of the CRISPR-Ca s system in executing immune functions.

A number of tools, including CRISPRTarget (Biswas et al., 2013), CASPERpam (Mendoza & Trinh, 2018), and Spacer2PAM (Rybnicky et al., 2022), have been developed to collect spacers from the CRISPR array and search the prokaryotic genome database for protospacer adjacent motifs. Spacer2PAM has demonstrated superior performance in guiding the experimental determination of functional PAMs in many instances; however, the quality and quantity of the input spacers are critical for its accuracy. When only a few spacers are available from the CRISPR array, the effectiveness of Spacer2PAM and similar tools may be significantly compromised. Moreover, Spacer2PAM requires the R package, taxonomizr, to convert alignment accession numbers into their corresponding genus and species names. The taxonomizr package necessitates the local downloading and organization of an SQL database, which occupies 65 GB of space.

In this work, we have developed PAMPHLET, a Python package designed to identify and guide the experimental determination of functional PAM sequences, especially for type I, II, and V CRISPR-Cas systems. This package enables the augmentation of the initial spacer input with a batch of spacers, allowing for a more stringent screening strategy in subsequent protospacer searches to reduce false positives. In comparison to SPACER2PAM, PAMPHLET demonstrated superior performance in most cases during our testing. This tool is particularly useful when the CRISPR array contains only a few spacers, as it utilizes the effector Cas protein sequence and a consistent repeat sequence as criteria to expand the original spacer input. Additionally, PAMPHLET does not require the local download and organization of an SQL database, thus saving significant storage space. Moreover, based on our precision tests on *Butyricimonas virosa* Cpf1, the predictions made by PAMPHLET are more stringent than those from *in vitro* tests by Doc MF, yet more consistent with *in vivo* results. In conclusion, PAMPHLET is a user-friendly and powerful tool for PAM prediction.

## 2. Materials and Methods

### 2.1 Prediction of PAM sequence

In the initial step, PAMPHLET searches for homologous proteins using either user-provided effector protein sequences or BLASTP (Camacho et al., 2009) results of effector proteins. A first protein threshold (FPT), defined as proident *≥*0.9 and procovs *≥* 0.9, is applied to filter potential homologous proteins. For those meeting the FPT criteria, we offer three selection modes for homologous proteins: ‘r’ (reference), ‘nr’ (non-reference), and ‘a’ (all). In ‘r’ mode, homologous proteins meeting the second protein threshold (SPT), with proident *≥* 0.98 and procovs *≥* 0.98, are considered as belonging to the reference species; other homologous proteins meeting the FPT and sharing species information with the reference species are also selected for further analysis. In ‘nr’ mode, we tally the species information of all homologous proteins meeting the FPT and select those belonging to the species with the highest frequency for subsequent analysis. In ‘a’ mode, we select all homologous proteins that meet the FPT without considering species information.

Next, the NCBI API is utilized to retrieve the source genomes of the selected homologous proteins. Once the genome sequences are obtained, MinCED (Bland et al., 2007) is used to identify all potential CRISPR arrays located within a *±*20kb range of the homologous protein positions in the genome. Following this putative CRISPR array prediction process, a pairwise alignment is generated between the consensus direct repeat from the predicted arrays and the user-provided consensus direct repeat. If the query coverage and percentage identity of the pairwise repeat alignment exceed the user-defined threshold (default: repcovs *≥* 0.9 and repident *≥* 0.9), the two consensus direct repeats are considered to be highly similar. Consequently, the CRISPR arrays containing these similar consensus direct repeats are deemed homologous. The distinct spacers from these homologous CRISPR arrays are then used to augment the original spacer dataset for subsequent analysis.

For the subsequent protospacer search step, two search modes are available: ‘pha-mode’ and ‘pro-mode’. ‘Pha-mode’ utilizes the spacers to search for protospacers in the BLASTN viruses database (taxid: 10239), while ‘pro-mode’ conducts searches in a non-eukaryotic database (excluding taxid: 2759). Additionally, two sets of parameters are offered for different BLASTN search strategies: ‘relax’ and ‘common’. The ‘common’ strategy uses the default parameters provided by NCBI BLASTN for proto- spacer discovery. The ‘relax’ strategy employs optimized parameters (WORD_SIZE=1, NUCL_REWARD=1, NUCL_PENALTY=-1, GAPCOSTS=5 2), which are particularly suited for spacers from type V arrays.

To screen for potential protospacers from BLASTN results, a strict threshold is applied. This requires that the alignment of protospacers must cover the entire spacer sequence with fewer than three mismatches and no gaps allowed. To address the issue of distinguishing mismatches or gaps at the ends of the spacer, the length of spacers that yield either no protospacer hits or only a single hit after screening will be adjusted. The primary method of length adjustment involves trimming one base from each end of the spacer. These shortened spacers will then undergo a second round of BLASTN searching. For subsequent analyses and calculations, protospacers originating from the same spacer are grouped together.

After completing the search for protospacer sources, the NCBI API will be utilized to download all relevant genome sequences. Subsequently, the sequences flanking each protospacer both upstream and downstream will be extracted. We will then calculate the score for each base at every position within each group of protospacers. Finally, scores for identical bases at corresponding positions across different groups will be aggregated to construct the position-base score matrix. This matrix is used to generate the logo plot. For this purpose, we use WebLogo 3.7.8 (Crooks et al., 2004) to create the logo plot and save the graphical result as a file.

### 2.2 DocMF PAM Prediction Screening

#### 2.2.1 Protein Purification

Cas genes were subcloned into the pET-28a vector to include an N-terminal 6*×*His tag. Both genes were expressed in the Escherichia coli BL21 (DE3) strain. After IPTG-induced expression and cell lysis, the Cas9 and Cas12a proteins were isolated using a HisTrapHP column (GE Healthcare) and stored in storage buffer (20 mM Tris-HCl pH 7.0, 150 mM NaCl, and 50% glycerol) at -80℃ for further research.

#### 2.2.2 gRNA Preparation

For Cas9 gRNA preparation, the double-stranded DNA (dsDNA) transcription templates were prepared by PCR amplification and then purified. For Cas12a crRNA preparation, a T7 promoter primer and crRNA_primer_R were mixed and incubated at 95°C for 3 minutes followed by 4°C for 10 minutes. These DNA products were then incubated with T7 RNA polymerase overnight at 37°C using the MEGAscript T7 Transcription Kit (Thermo Fisher Scientific), and the RNA was purified using the GeneJET RNA Cleanup and Concentration Micro Kit (Thermo Fisher Scientific). RNA concentration was quantified using the Qubit RNA High Sensitivity Assay Kit (Thermo Fisher Scientific) and measured with a Qubit Fluorometer. The sequences of the primers are listed in Supplementary File S1.

#### 2.2.3 DocMF Experiments

For PAM identification experiments, the sequence of the single-stranded DNA (ssDNA) oligonucleotide library used to create single-stranded circles is detailed in Table S1 and was supplied by BGI TECH SOLUTIONS (BEIJING LIUHE) CO., LIMITED. A quantity of 0.5 pmol of ssDNA was used for Single-stranded Circle (ssCir) and DNA Nanoball (DNB) preparation as per the instructions in the BGISEQ-500RS High-throughput Sequencing Kit (SE100). The concentration of DNBs was quantified using the Qubit ssDNA Assay Kit (Thermo Fisher Scientific). PAM identification was carried out according to the method described in Li et al., 2020, with a minor modification: 100 µL of DNBs were loaded onto the BGISEQ-500 V3.1 chip using 33 µL of BGISEQ-500 DNB Loader (MGI Tech Co. Ltd.). Single-end sequencing runs for 43 bases were conducted following the BGISEQ-500RS High-throughput Sequencing Kit instructions.

### 2.3 Cell Culture and Transfection

Add 10% fetal bovine serum and 1% non-essential amino acids to Dulbecco’s Modified Eagle’s Medium when cultivating 293T cells. When the cell density reaches 80%–90%, use TrypLE™ Express for digestion and passage. The day before transfection, seed the cells onto a 12-well plate at a density of 2 *×* 10^5^ cells per well. When confluency reaches approximately 50%, transfect the cells with plasmids using Lipofectamine™ 3000 according to the manufacturer’s instructions, adding 1000 ng of plasmid DNA per well. Observe the transfection efficiency by fluorescence analysis 48–72 hours post-transfection.

### 2.4 Indel Detection Assays

After calculating the fluorescence rate, collect the cells and extract genomic DNA using Tiangen’s reagent kit. Amplify the region of interest through PCR using PrimeStar GXL (Takara) with 100 ng of genomic DNA as the template. Use the following primers: PCR Forward Primer: 5’-TTTCCGGAGCACTTCCTTCT-3’ and PCR Reverse Primer: 5’-CAGGAACCCCTGTAGGGAAG-3’. Purify the PCR product using the NucleoSpin Gel and PCR Clean-up Kit, following the manufacturer’s instructions. Denature and reanneal 50 ng of the purified PCR product, followed by digestion with 0.3 µL of T7E1 (New England Biolabs) at 37°C for 20 minutes. After the reaction, add 10*×* loading buffer and perform agarose gel electrophoresis.

### 2.5 CRISPR-Cas system prediction and co-occurrence

The data for this study were sourced from the comprehensive prokaryotic genome assemblies available in the National Center for Biotechnology Information (NCBI) database (Sayers et al., 2021) as of the specified reference date, 09/11/2023. We employed the CRISPRCasTyper (Russel et al., 2020) tool to perform predictive analyses of CRISPR-Cas systems within the prokaryotic genomic data. Each identified CRISPR-Cas system was rigorously evaluated for the completeness of its cas operon. We selectively excluded Class I CRISPR-Cas systems that presented with an incomplete adaptation module, a critical component required for the spacer acquisition process. For the purposes of this study, co-occurring CRISPR-Cas systems were defined as different subtypes residing within the same genomic assembly. These were annotated as co-occurrences within our dataset. Conversely, CRISPR-Cas systems containing either a single subtype or several occurrences of the same subtype within one assembly were categorized as independent systems.

## 3. Results

### 3.1 Design of PAMPHLET

PAMPHLET is designed to predict potential Protospacer Adjacent Motifs (PAMs) for a given CRISPR-Cas system. It requires the sequences of Cas proteins, CRISPR array spacers, and the consensus repeat sequence of the array as inputs. The tool is comprised of three main modules: 1) spacer expansion; 2) protospacer identification; and 3) PAM prediction (Figure 2).

**Figure 2:**
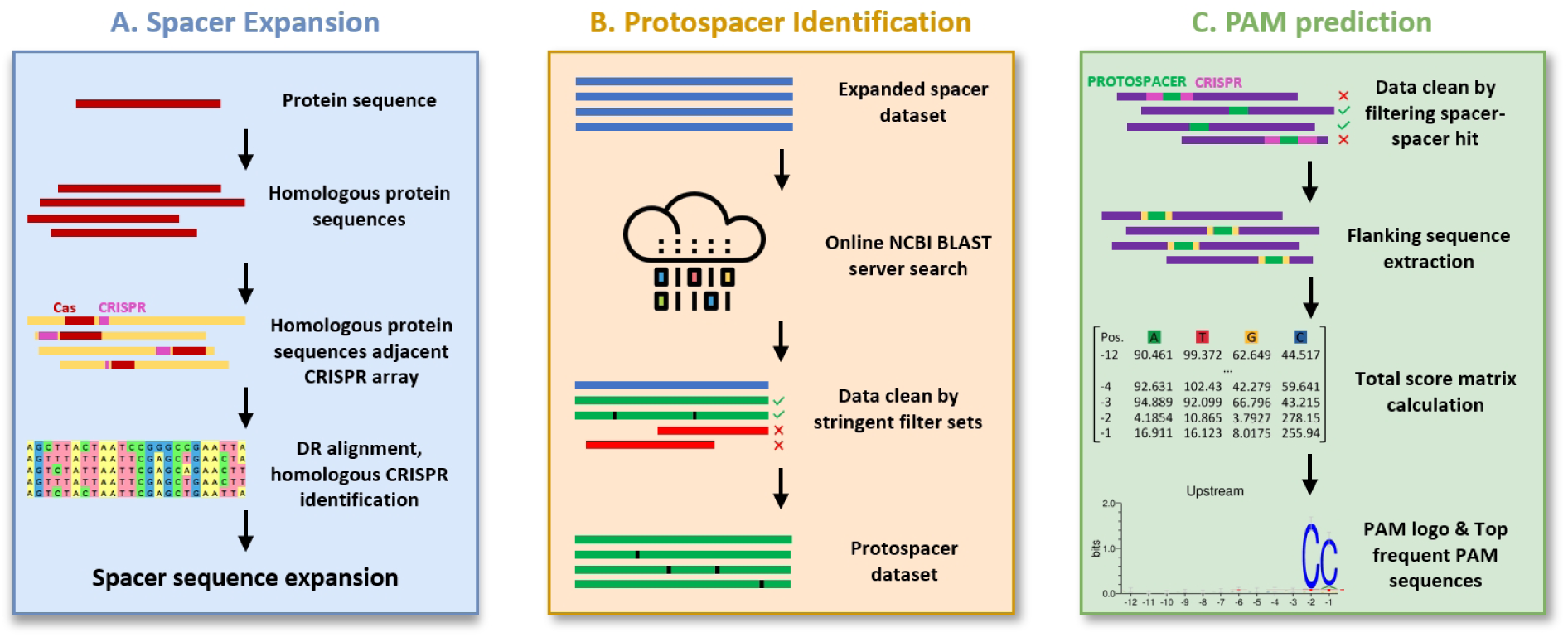
A concise overview of the PAMPHLET workflow, which is composed of three primary modules: **(A)** spacer expansion; **(B)** protospacer identification; and **(C)** PAM prediction

The primary function of the spacer expansion module is to broaden the spacer database—that is, to identify additional spacers for PAM prediction. To achieve this, we developed an innovative method to discover homologous CRISPR-Cas systems that exhibit high similarity to the input system. According to our criteria, CRISPR-Cas systems are considered homologous if their Cas proteins and Direct Repeat (DR) sequences show at least 90% sequence similarity to those of the target CRISPR-Cas system. Systems that are homologous are expected to have identical PAMs because of their high degree of similarity. The process used to identify homologous CRISPR-Cas systems and expand the spacer database is depicted in Figure 1A and includes the following steps:

Step 1. We performed a search for the input Cas proteins against the NCBI nr database to find homologous Cas proteins in other genomes, with similarity defined as 90% or higher.

Step 2. We predicted and identified CRISPR arrays in proximity to the homologous Cas proteins discovered in Step 1.

Step 3. We compared the consensus repeat sequences of the CRISPR arrays from Step 2 to that of the input CRISPR-Cas system. Only those systems with a consensus repeat sequence similarity of 90% or greater were considered homologous and included in the spacer database.

Spacers from the homologous CRISPR-Cas systems will be added to our spacer database and used along with the spacers from the provided CRISPR-Cas system to identify potential protospacers.

In the protospacer identification module (see Figure 2B), spacers identified in the previous module are used as queries and automatically submitted to the NCBI BLAST server online. Stringent filters (qcov = 0.9, evalue *≤* 1) are applied to the resulting hit table to select alignments that are likely to represent protospacers.

These selected alignments are then potentially used to determine the genomes that harbor the putative protospacers and to predict the likely locations of PAMs.

In developing the PAM prediction module, illustrated in Figure 2C, we implemented a filtering protocol to remove candidate protospacers located within CRISPR-Cas loci. This step is essential to prevent the incorrect identification of CRISPR array repeats as PAM sequences and to maintain the accuracy of PAM identification.

Following this refinement, we carefully extracted 12-bp sequences from both sides of the cleaned protospacer group, obtaining upstream and downstream sequences crucial for PAM prediction. Although the default setting is a 12-bp length, our tool allows users to define custom lengths for these flanking regions to meet various research needs.

To aid computational analyses, we have established a positional nomenclature for the 12-bp flanking sequences: upstream segments are labeled as [-12, -11, …, -2, -1], and downstream segments as [+1, +2, …, +11, +12]. For a specific spacer, represented by *spacer_i_*, we calculate a frequency matrix *F* (*spacer_i_*) based on the flanking sequences of its corresponding protospacers. This matrix is a critical component for statistically modeling PAM sequence prediction, utilizing the valuable information contained within these regions.

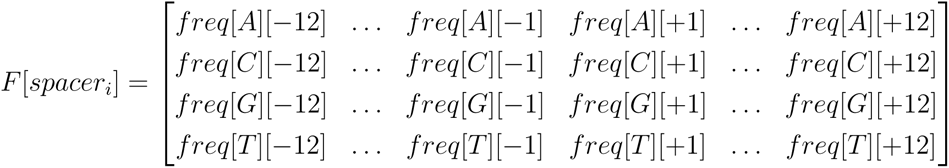

The calculation of *freq*[*base*][*pos*], which represents the frequency of nucleotides A, T, C, or G at a specific position, is as follows:

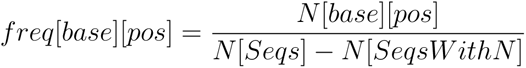

To mitigate potential bias caused by varying counts of flanking sequences across different spacers, which may skew analytical results, our algorithm PAMPHLET employs a logistic normalization function. This method systematically adjusts the raw frequency matrix *F* (*spacer_i_*) to account for size disparities, thus producing a calibrated and weighted score matrix *S*[*spacer_i_*] for each spacer:

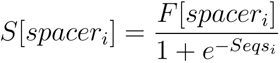

where *Seqs_i_*is the total number of flanking sequences for *spacer_i_*.

In the final phase of analysis, we combine the individual weighted score matrices *S*[*spacer_i_*] for each spacer into a cumulative score matrix *T*. This matrix is formed by summing the weighted contributions from all the spacers being studied. PAMPHLET then uses this consolidated matrix *T* to generate a WebLogo plot:

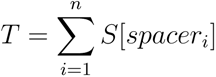

where *n* represents the total number of spacers, including both original and expanded spacers, PAMPHLET not only generates a WebLogo plot but also provides a table that lists all flanking sequences, sorted by their frequency of occurrence.

### 3.2 Assessing PAMPHLET’s Predictive Precision and Expansion Efficacy in Diverse CRISPR-Cas Systems

Here, we evaluate PAMPHLET’s performance on a range of CRISPR-Cas systems with experimentally validated PAMs derived from the following species: *Acinetobacter baumannii* (Karah et al., 2015), *Bacillus halodurans* (Leenay et al., 2016), *Campylobacter jejuni* (Kim et al., 2017), *Clostridioides difficile* (Boudry et al., 2015), *Clostridium pasteurianum* (Pyne et al., 2016), *Clostridium tyrobutyricum* (Zhang et al., 2018), *Francisella tularensis* (Zetsche et al., 2015), *Gluconobacter oxydans* (Qin et al., 2021), *Hungateiclostridium thermocellum* (Walker et al., 2020), *Lactobacillus cripatus* (Hidalgo-Cantabrana et al., 2019), *Moraxella bovoculi* (Zetsche et al., 2015), *Neisseria meningitidis* (Esvelt et al., 2013), *Parvibaculum lavamentivorans* (Ran et al., 2015), *Pseudomonas aeruginosa* (Cady et al., 2012), *Staphylococcus aureus* (Ran et al., 2015), *Streptococcus canis* (Chatterjee et al., 2018), *Streptococcus pasteurianus* (Ran et al., 2015), *Streptococcus pyogenes* (Jinek et al., 2012), *Streptococcus thermophilus* (Garneau et al., 2010), and *Treponema denticola* (Esvelt et al., 2013). With PAMPHLET, the number of spacers expanded ranges from 1 to 18.25 times (see Figure 3A). We compare these results with those from Spacer2PAM; PAMPHLET accurately predicts 18 out of 20 systems either perfectly or functionally, and it outperforms Spacer2PAM in 9 out of 18 comparisons (see Figure 3C and Supplementary File S2).

**Figure 3:**
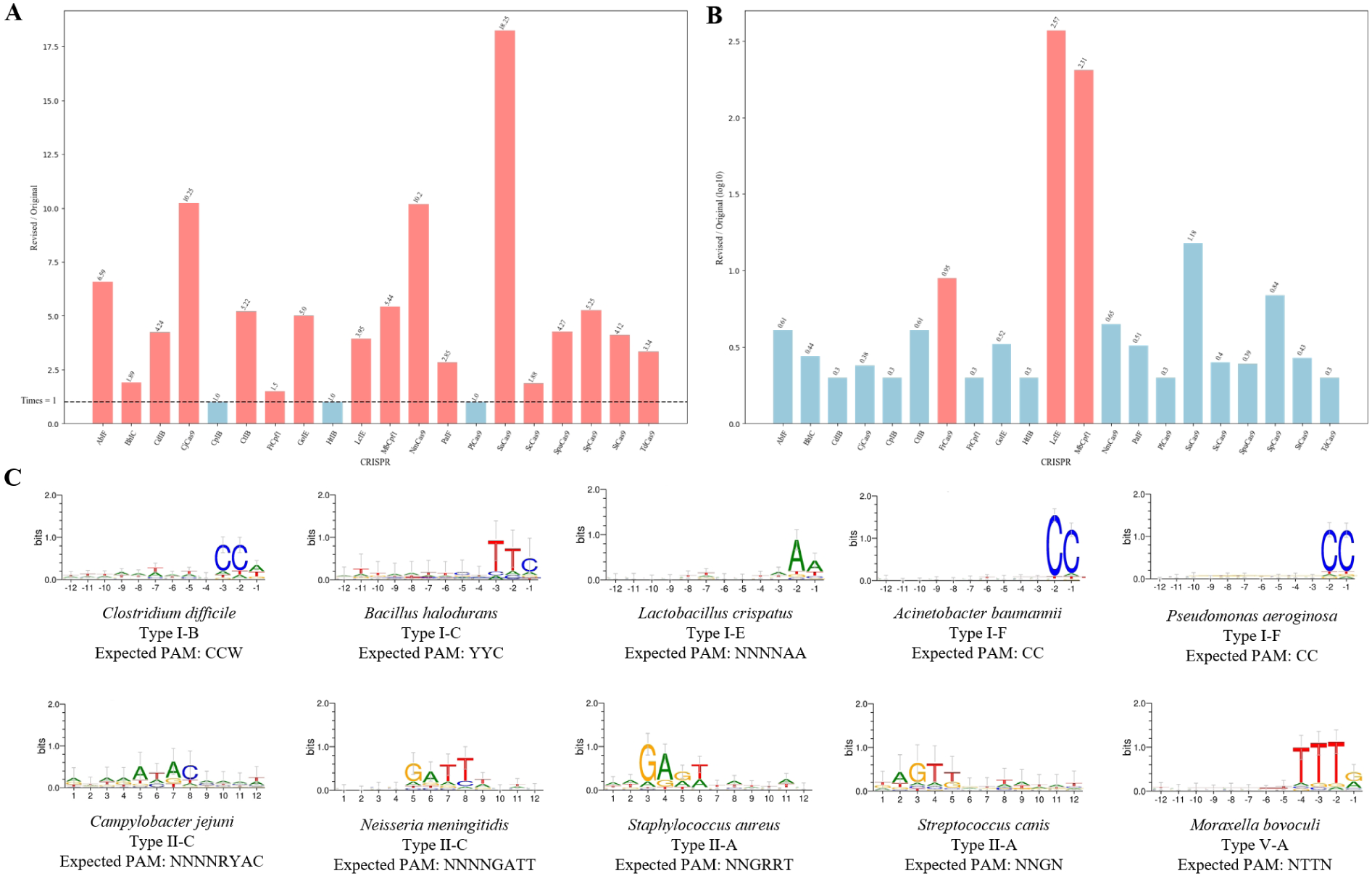
PAM prediction evaluation of PAMPHLET. **(A)** The ratio of the original spacer number to the expanded spacer number; **(B)** Ratio of protospacers retrievable by original spacers to those retrievable by expanded spacers; **(C)** A selection of PAM prediction results.

To further demonstrate the homology extension algorithm’s effectiveness in identifying viable protospacers, we calculated the ratio of protospacers detectable by original spacers against those by expanded spacers. Within our dataset, we found that while some candidates’ original spacers failed to match any protospacers, others showed a marked increase in the detection of viable protospacers, as depicted in Figure 3B. These results underscore the utility of homology-based extension strategies in augmenting the pool of effective protospacers, which in turn enhances the accuracy of PAM sequence predictions. The improved performance suggests that expanded spacers can effectively close the gap in recognition between spacers and protospacers, thus boosting PAM prediction efficiency.

### 3.3 Wet-lab validation of PAMPHLET

#### 3.3.1 Enhanced Specificity and Nuanced PAM Recognition by PAMPHLET Compared to DocMF

To further validate the predictive accuracy of the PAMPHLET algorithm, we performed a detailed comparative analysis of protospacer adjacent motif (PAM) requirements for both type II-A and type V-A CRISPR-Cas systems, using DocMF for comparison. We examined the PAM consensus sequences for the *Veillonella* sp. Cas9, as shown in Figure 4. The sequences derived from PAMPHLET were identified as 5’-NTARRNN-3’, indicating a more distinct nucleotide preference at the second position compared to those identified by DocMF, which were 5’-NNAAANN-3’. This increased specificity suggests that PAMPHLET may more accurately discern the stringent recognition patterns of PAMs, potentially capturing subtle distinctions between *in vivo* and *in vitro* PAM requirements. For the *Segatella copri* Cpf1, the PAMPHLET algorithm accurately identified the PAM sequence as 5’-TTTN-3’, revealing significant conservation of thymine residues at the -3 and -4 positions—a specificity surpassing that provided by DocMF. This notable difference may stem from the varying sensitivity to PAM recognition in the DocMF method, which, under *in vitro* cleavage conditions, might become less stringent due to enzyme overabundance, leading to broader PAM recognition.

**Figure 4:**
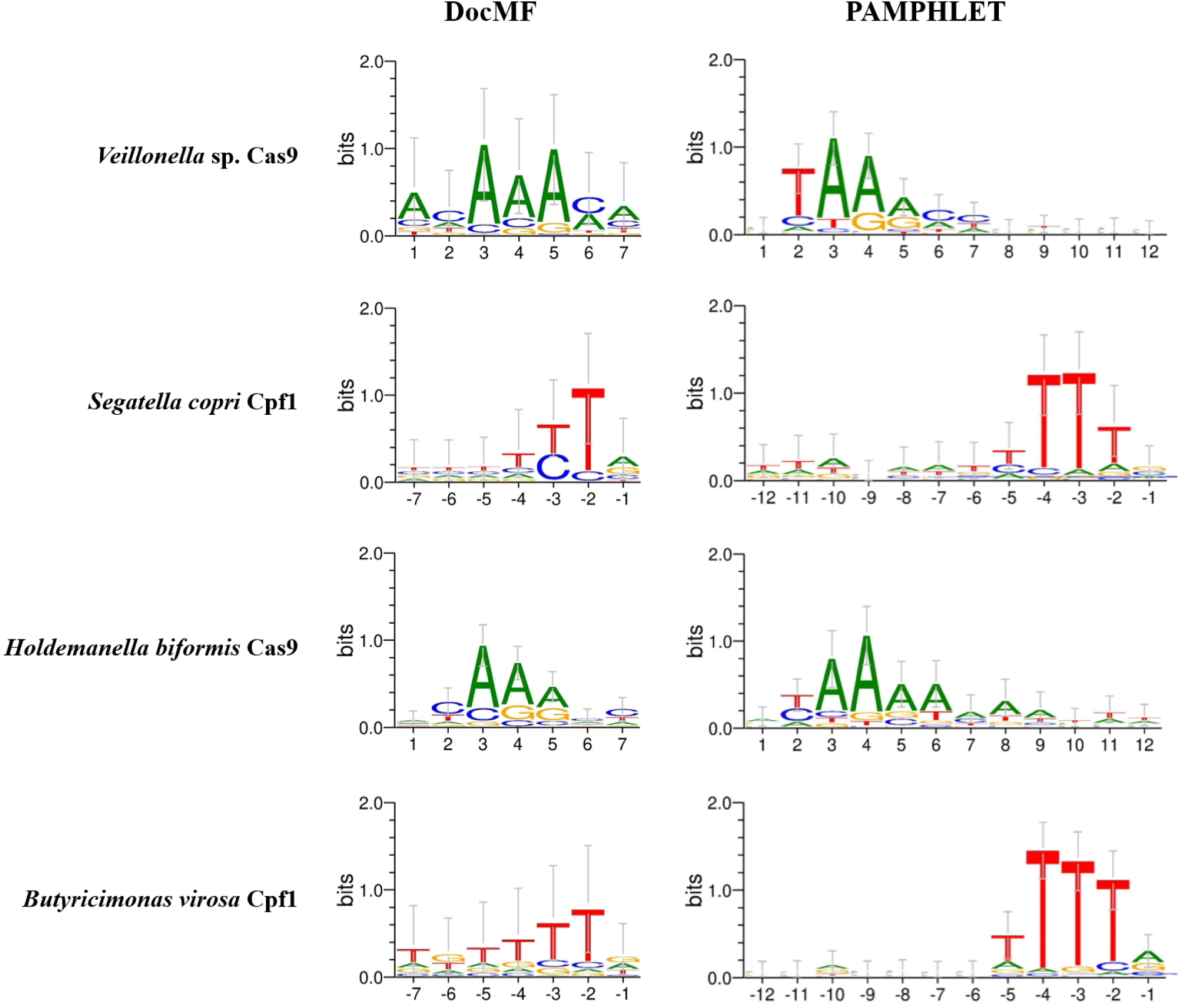
PAM provided by pamphlet versus PAM verified by DocMF.

Parallel findings were obtained for *Holdemanella biformis* Cas9 and *Butyricimonas virosa* Cpf1. In summary, the application of PAMPHLET confers an enhanced level of precision in delineating stringent PAM sequences, thereby affirming its utility as a superior tool for defining the molecular determinants of PAM specificity in CRISPR-Cas systems. In the case of *Holdemanella biformis* Cas9, the predictive outcomes highlight a distinct conservation bias for adenine residues at the +3 and +4 positions. While DocMF predictions indicated a preference for adenine conservation at the +3 position, PAMPHLET revealed a pronounced preference at the +4 position. This contrast in sequence conservation underscores the advanced predictive capabilities of PAMPHLET in identifying position-specific nucleotide preferences that may not be as evident in the profiles generated by DocMF.

Likewise, for *Butyricimonas virosa* Cpf1, a comparative analysis between DocMF and PAMPHLET predictions indicated a gradient in the conservation of thymine residues from the -2 to -4 positions. DocMF’s predictions suggested a decreasing emphasis on thymine conservation, with a significant reduction at the -4 position, which was considered non-essential. In stark contrast, PAMPHLET’s results assigned equal importance to the thymine at the -4 position, challenging the gradient of conservation proposed by DocMF and underscoring the significance of this nucleotide in the PAM sequence context.

#### 3.3.2 Corroborating PAMPHLET’s PAM Predictions with Targeted Genome Editing in HEK293T Cells

The observed discrepancy in the PAM specificity of *Butyricimonas virosa* Cpf1, especially concerning the conservation of thymine at the -4 position, prompted us to reconcile computational predictions with biological reality. PAMPHLET’s algorithm suggested thymine at this position is indispensable, in contrast to the more permissive prediction from DocMF.

To empirically validate the predictive output of our PAMPHLET algorithm, we engineered plasmids encoding guide RNAs (gRNAs) alongside *Butyricimonas virosa* Cpf1 nuclease and conducted a series of genome editing assays in HEK293T cell populations. The plasmids were co-labeled with a green fluorescent protein (GFP) reporter to facilitate monitoring of transfection efficiency. Negative control plasmids (referred to as NC plasmids) were designed to express a non-functional gRNA. Targeting the *AAVS1* genomic locus, we selected six distinct genomic sites (listed in Table 1), close enough to allow the use of uniform primer sets for subsequent PCR amplification. The cleavage efficacy of *Butyricimonas virosa* Cpf1 was quantitatively assessed through T7 endonuclease I (T7E1) mismatch detection assays. Results shown in Figure 5 indicate pronounced enzymatic activity for *Butyricimonas virosa* Cpf1 with a 5’-TTTN PAM sequence, while alternative PAM configurations showed negligible activity. This confirms a high specificity for PAM recognition by *Butyricimonas virosa* Cpf1, in line with our initial predictions from PAMPHLET, thus supporting the reliability of our bioinformatic tool in precisely predicting PAM constraints in a cellular context.

**Figure 5:**
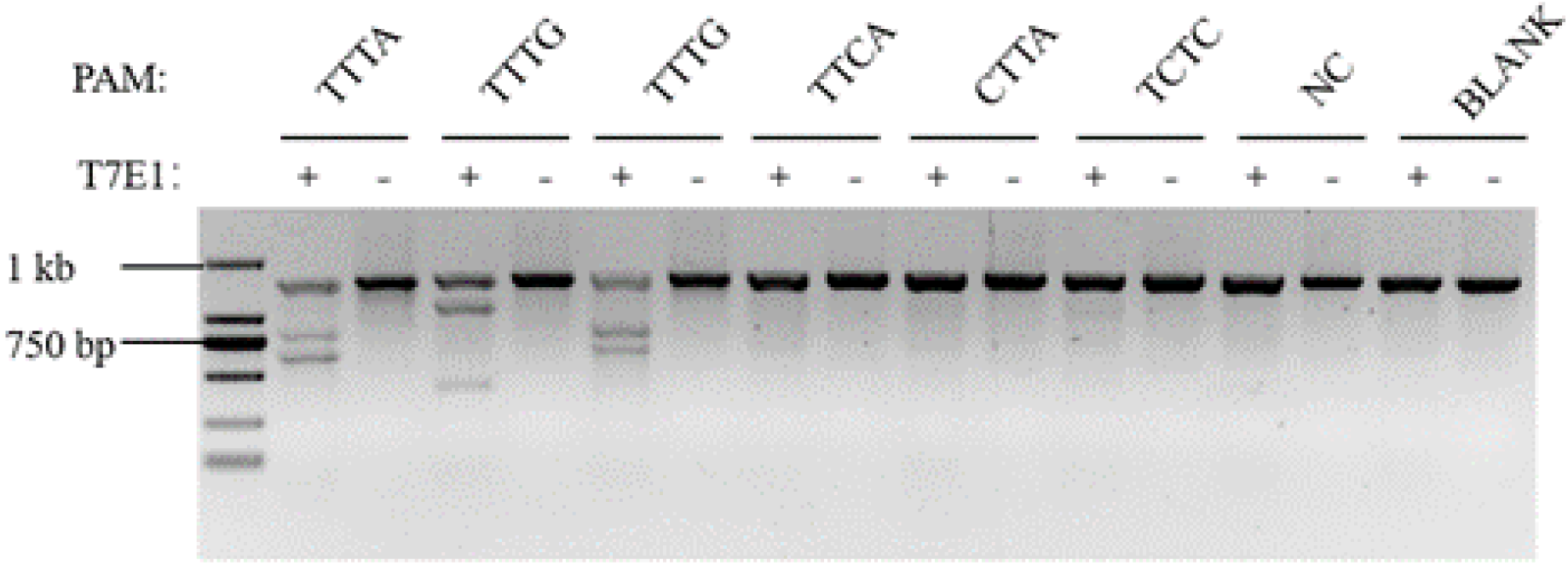
Genome editing in human HEK293T cells using BvCas12a.

**Table 1:**
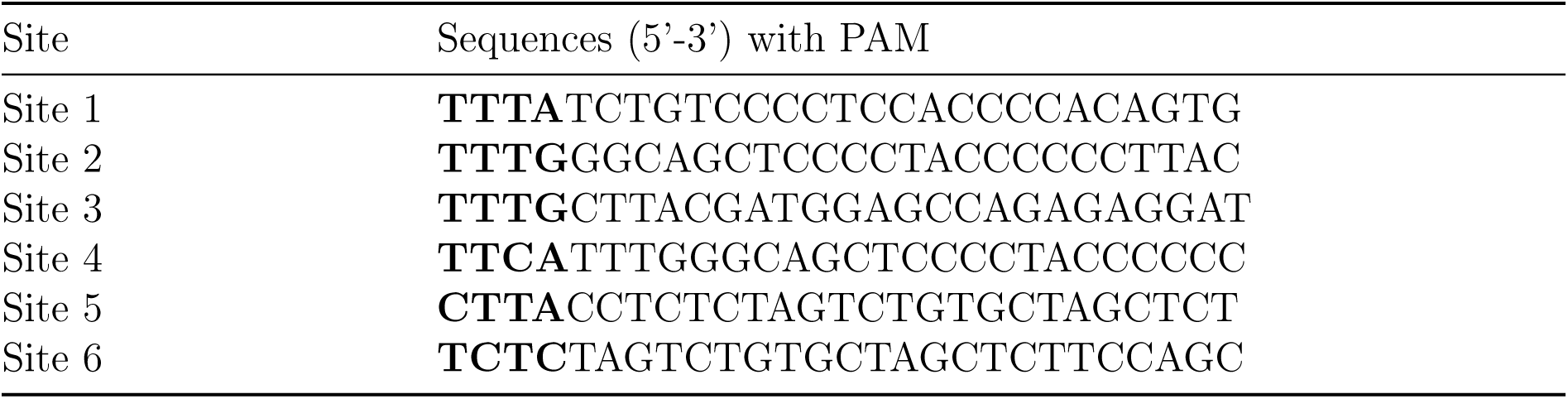
Target sequences of *Butyricimonas virosa* Cpf1 at *AAVS1*.

### 3.4 co-occurrence systems

Within prokaryotic genomes, the co-occurrence of multiple distinct CRISPR-Cas systems has been confirmed. These systems likely represent a plethora of adaptive immune mechanisms employed by prokaryotes to fend off exogenous genetic elements, such as bacteriophages and plasmids. Here, we conducted a predictive analysis and co-occurrence statistics of the CRISPR-Cas systems within the prokaryotic genomes available in the National Center for Biotechnology Information (NCBI) database.

Upon analyzing the dataset comprising 311,860 prokaryotic genomes, we utilized bioinformatics tools to predict the presence of CRISPR-Cas systems. We identified a total of 85,353 bacterial strains encoding CRISPR-Cas systems, collectively containing 97,997 CRISPR-Cas systems. These systems are representative of the entire currently known spectrum of CRISPR-Cas system families. Notably, within these identified systems, there was a higher representation of types I, II, and III CRISPR-Cas systems. This distribution reflects the prevalence and potential importance of these CRISPR-Cas systems in prokaryotic immunity. In stark contrast, Class V CRISPR-Cas systems, particularly those akin to the mini-Cas variants such as Cas12f1 and Cas12n, manifested as a minority, with a mere 0.10% representation within the dataset.

In our subsequent classification, 62,157 strains were identified that encode a single CRISPR-Cas system, while 2,689 strains harbored multiple, distinct CRISPR-Cas systems. Our statistical analysis on the co-occurrence of Class I and Class II systems, with attention to the diversity of effector modules, revealed that type II-A systems frequently co-occur with other systems, likely due to their abundance. Interestingly, the rarer type V-F systems were also found to co-occur notably with type I-B systems (Figure 6). The absence of an adaptation module, comprised of Cas1 and Cas2 proteins, in the V-F1 system piqued our interest in its spacer acquisition mechanism. We hypothesize that V-F1 may leverage the adaptation module of a co-existing CRISPR system for new spacer integration. If this hypothesis is correct, we anticipate significant homology in the protospacer adjacent motifs (PAMs) between the V-F1 system and its hypothetical donor system.

**Figure 6:**
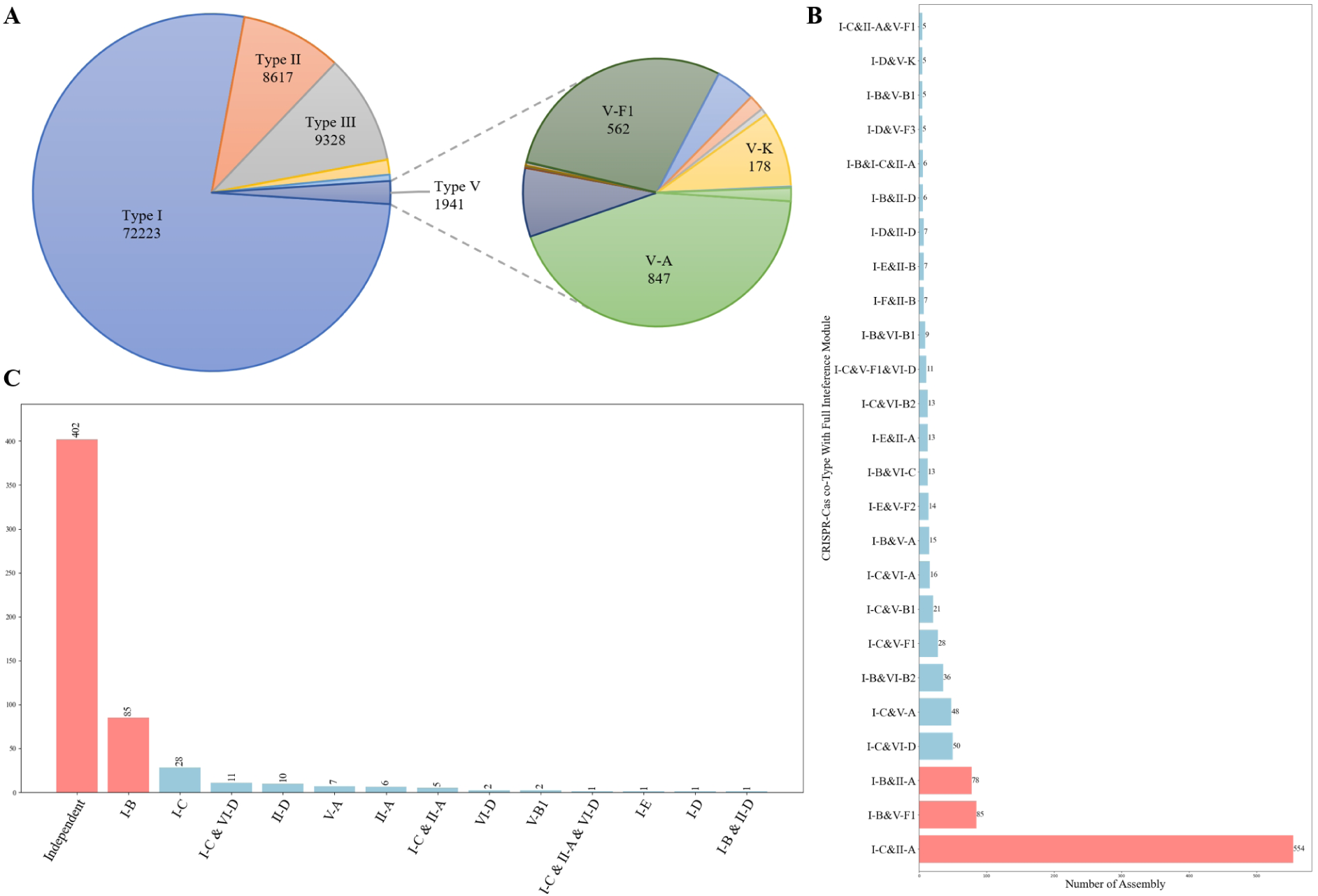
Comprehensive Statistical Analysis of CRISPR-Cas Systems in Prokaryotic Genomes. **(A)** Typing and Subtyping of CRISPR-Cas Systems with Emphasis on Type V Variants; **(B)** Co-occurrence Statistics of CRISPR-Cas Systems; **(C)** Independent Versus co-occurrence Type V CRISPR-Cas Systems Statistics.

To test our hypothesis, we analyzed PAM sequences of V-F subtype CRISPR-Cas systems that operate independently as well as those co-occurring with I-B subtype systems. Detailed results of this predictive analysis are available in Supplementary File S3 and S4. Notably, the PAMPHLET tool showed limited effectiveness in identifying PAMs for certain V-F type CRISPR-Cas systems. Of the 85 co-occurring systems we examined, only 26 enabled the accurate prediction of both I-B and V-F PAMs. Further, we found that 14 of these co-occurring systems shared PAM sequences, a finding illustrated in Figure 7.

**Figure 7:**
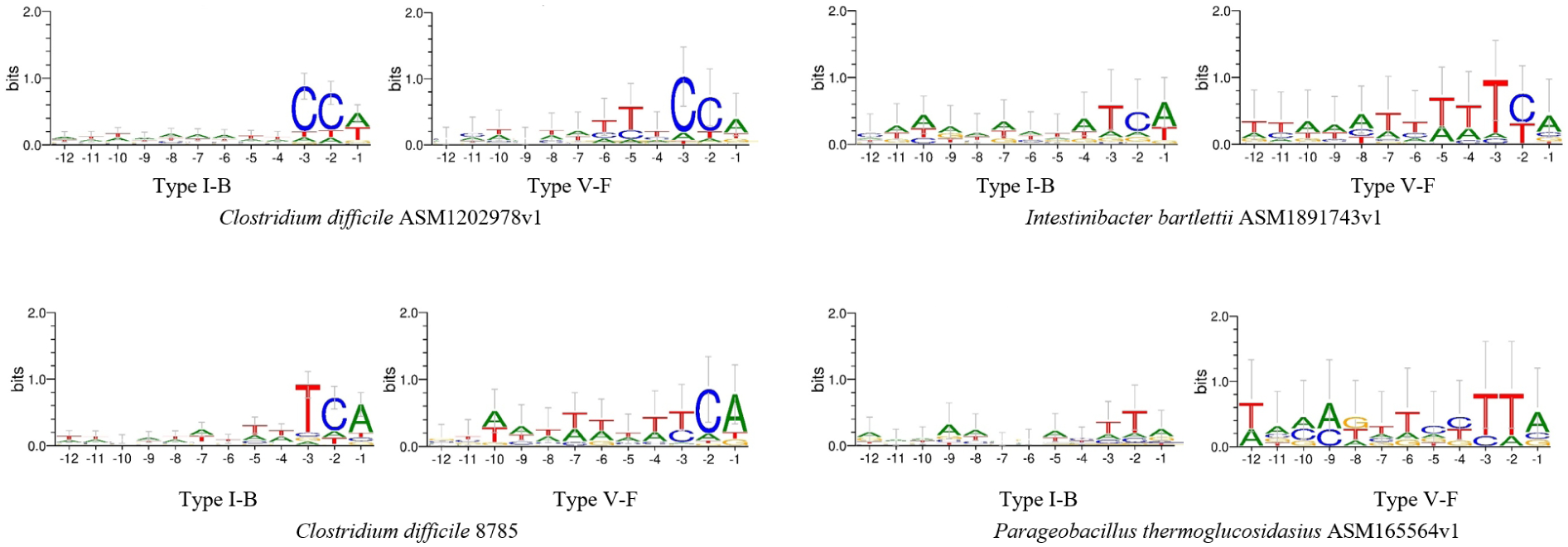
Representative PAM for subtype V-F systems co-occurrence with the subtype I-B.

The increased similarity of PAM sequences within co-occurring systems poses a challenging puzzle that defies a simple biological explanation. Although we suspect that a system might use the adaptation module from a co-existing CRISPR system to incorporate new spacers, we cannot yet experimentally confirm this theory. If our theory is proven, it would suggest significant homology in the PAMs between such co-existing systems, notably between the V-F1 system and its proposed donor. However, due to current experimental limitations, our idea remains conjectural, necessitating further research to elucidate the mechanism and implications of this PAM sequence similarity. Unraveling this mystery will undoubtedly deepen our comprehension of the evolutionary and functional dynamics of CRISPR-Cas systems.

## 4. Discussion

In this work, we established PAMPHLET using only five Python packages. The tool incorporates a Homologue-Enhanced Strategy to augment the spacer dataset, thereby addressing the issue of obtaining meaningless results from SPACER2PAM due to a scarcity of spacers. Additionally, PAMPHLET’s predictive outputs often outperformed those of SPACER2PAM and demonstrated high consistency with both DocMF and in vivo validation.

The default parameters for the Homologue-Enhanced Strategy are set to be highly stringent, creating a paradox between the quality and the quantity of information it yields. On one hand, we posit that proteins, particularly those in the PI domain with high sequence similarity, will recognize identical PAMs. Such similarity within proteins and direct repeats (DRs) implies greater confidence in the results. On the other hand, we aim to effectively expand the spacer datasets to gather ample protospacer and adjacent sequence information. Although our default parameters have been tested across various subtypes and have shown promising performance in most cases, users have the flexibility to adjust these parameters to find an optimal balance between the quality and quantity of the resulting information.

We provided users with the option to use either a prokaryote or phage database for protospacer online BLASTN searches. Initially, we posited that spacers in most CRISPR arrays originated from bacteriophages. Accordingly, we employed a viral database (taxid:10239) for BLASTN searches, which yielded successful results for many subtypes. However, in certain subtypes, the phage database failed to return expected protospacer hits, resulting in null results. We attributed these null results to the limited size of the phage database. Substituting the phage database with a prokaryote database (excluding taxid:2759) decreased the incidence of null results. Upon analyzing the bacterial genomes where these protospacer hits were found, we observed that the hits were primarily located in CRISPR arrays, inserted prophage regions, and plasmids, corroborating our hypothesis of bacteriophage derivation. This insight may provide a new avenue for identifying potential prophages.

The PAM we predicted may be more stringent than actual ones. Our tool operates on the premise that proteins involved in both the adaption and interference stages of the same CRISPR system recognize identical PAMs. Effector proteins are necessary for spacer acquisition in some systems (Heler et al., 2015). However, in many cases, these proteins are not part of the adaptation process (Xiao et al., 2017). Despite this, the PAMs identified by proteins involved in adaptation are also recognized by effector proteins, suggesting a coevolutionary relationship between adaptation and interference (J. Wang et al., 2015). We hypothesize that effector proteins in interference stages generally recognize a broader range of PAMs compared to proteins in adaptation, to ensure the effectiveness of all captured spacers in the defense process. Furthermore, research indicates that there is an arms race between bacterial defense systems and phages. One strategy for phages to evade CRISPR-Cas immunity is through high-frequency mutations in PAM and seed regions (Common et al., 2019; Vale & Little, 2010). Therefore, we propose that a more relaxed PAM recognition by bacteria enhances their ability to counteract the rapid mutation rates found in phage genomes.

The PAM we predicted may exhibit divergences from the DocMF in certain instances. There is a debate over whether the PAM derived from *in vitro* experiments is entirely consistent with the functional PAM that effector proteins genuinely recognize *in vivo*. For example, the PAM we predicted for the type V-A system in *Moraxella bovoculi* indicates a preference for a T or C base at the -4 position upstream, a preference not observed in *in vitro* PAM screening experiments. In practice, *in vivo* testing revealed a significant C/T preference at this position, as evidenced by testing four guides per NTTN PAM *in vivo*, each targeting a different gene (*DNMT1*, *EMX1*, *GRIN2b*, or *VEGFA*). Furthermore, *in vivo* PAM testing on *Butyricimonas virosa* Cpf1 confirmed a stringent 5’TTTN PAM, which aligned with our predictions, while the DocMF suggested that the -4 position was of low importance. This discrepancy may be due to varying ratios of Cas enzyme to target DNA in different conditions. In conclusion, our tool demonstrates improved accuracy over *in vitro* PAM screening methods.

PAMPHLET still has some limitations. First, the ineffective expansion of spacer datasets diminishes PAMPHLET’s performance. If only a few homologous proteins are found, there will be minimal expansion of the spacer datasets, and PAMPHLET’s performance will be similar to that of SPACER2PAM. Second, some false positive self-hits are inevitable. During the protospacer search, spacers from other CRISPR arrays can be mistakenly identified as targets. Although we have employed MinCED to call CRISPR arrays as a checkpoint to filter out such self-targeting hits, it fails to identify some orphan CRISPR arrays of low quality. Third, MinCED’s limitations can lead to the misidentification of spacers. Canonical CRISPR arrays consist of identical DRs and diverse spacers. However, ‘half DRs,’ which include only part of a DR, are often found at the ends of CRISPR arrays. MinCED struggles to determine the correct boundaries of DRs and repeats, sometimes incorrectly truncating the DR region to match these ‘half DRs’ and mistakenly assigning parts of DRs to spacers, resulting in impure spacers that are discarded in subsequent steps. Lastly, PAMPHLET performs poorly with some type V systems. For instance, PAMPHLET yielded no results for V-J systems from bacteriophages, where the sequence identity between any two characterized CasΦ proteins is approximately 30-40%. This low level of similarity means that the homology-enhanced strategy is ineffective for this protein family.

As more Type V subtypes lacking Cas1 and Cas2 are characterized, the mechanisms underlying the formation of their CRISPR arrays are not yet fully understood. We find it particularly intriguing that subtype V-F1 has a significantly higher co-occurrence ratio with subtype I-B. Indeed, previous research has suggested that Type III systems, which lack certain Cas proteins for spacer acquisition and sometimes even a CRISPR array, might utilize the CRISPR array of Type I. We hypothesize that V-F1 may adopt the spacer acquisition mechanism of the I-B array, by providing evidence of their PAM consistency via PAMPHLET predictions. Given that the Cas1,2 complex captures heterologous DNA and recognizes and cuts repeats in the array to a fixed length, the length of newly inserted spacers and repeats is consistent with the existing ones. The similarities between V-F1 and I-B could shed light on how the Cas1,2 complex recognizes repeats, offering insights into the evolutionary trajectory of Type V CRISPR-Cas systems from IS transposases with TnpB.(Abdelrahman et al., 2021)

In conclusion, PAMPHLET introduces a novel homology-enhanced strategy that significantly improves performance. Despite some remaining limitations, it proves to be extremely valuable for determining PAM sequences in novel CRISPR-Cas systems.

## 5. Code Availability

Source code is available at https://github.com/GolshkovQ/PAMPHLET

## 6. Acknowledge

The authors declare that the research was conducted in the absence of any commercial or financial relationships that could be construed as a potential conflict of interest. The authors wish to thank the China National GeneBank (Shenzhen) for their support in our research. We are also grateful to BNU-HKBU UIC for providing the necessary academic resources and environment that greatly contributed to this study. Our acknowledgment also extends to the funding bodies associated with BNU-HKBU UIC for their financial support.

## References

Abdelrahman, M., Wei, Z., Rohila, J. S., & Zhao, K. (2021). Multiplex genome-editing technologies for revolutionizing plant biology and crop improvement. Front. Plant Sci., 12, 721203.

Armario Najera, V., Twyman, R. M., Christou, P., & Zhu, C. (2019). Applications of multiplex genome editing in higher plants. Curr. Opin. Biotechnol., 59, 93–102.

Barrangou, R., Fremaux, C., Deveau, H., Richards, M., Boyaval, P., Moineau, S., Romero, D. A., & Horvath, P. (2007). CRISPR provides acquired resistance against viruses in prokaryotes. Science, 315(5819), 1709–1712.

Biswas, A., Gagnon, J. N., Brouns, S. J. J., Fineran, P. C., & Brown, C. M. (2013). CRISPRTarget: Bioinformatic prediction and analysis of crRNA targets. RNA Biol., 10(5), 817–827.

Bland, C., Ramsey, T. L., Sabree, F., Lowe, M., Brown, K., Kyrpides, N. C., & Hugenholtz, P. (2007). CRISPR recognition tool (CRT): A tool for automatic detection of clustered regularly interspaced palindromic repeats. BMC Bioinformatics, 8(1), 209.

Boudry, P., Semenova, E., Monot, M., Datsenko, K. A., Lopatina, A., Sekulovic, O., Ospina-Bedoya, M., Fortier, L.-C., Severinov, K., Dupuy, B., & Soutourina, O. (2015). Function of the CRISPR-Cas system of the human pathogen clostridium difficile. MBio, 6(5), e01112–15.

Briner, A. E., Donohoue, P. D., Gomaa, A. A., Selle, K., Slorach, E. M., Nye, C. H., Haurwitz, R. E., Beisel, C. L., May, A. P., & Barrangou, R. (2014). Guide RNA functional modules direct cas9 activity and orthogonality. Mol. Cell, 56(2), 333– 339.

Cady, K. C., Bondy-Denomy, J., Heussler, G. E., Davidson, A. R., & O’Toole, G. A. (2012). The CRISPR/Cas adaptive immune system of pseudomonas aeruginosa mediates resistance to naturally occurring and engineered phages. J. Bacteriol., 194(21), 5728–5738.

Camacho, C., Coulouris, G., Avagyan, V., Ma, N., Papadopoulos, J., Bealer, K., & Madden, T. L. (2009). BLAST+: Architecture and applications. BMC Bioinformatics, 10(1), 421.

Chatterjee, P., Jakimo, N., & Jacobson, J. M. (2018). Minimal PAM specificity of a highly similar SpCas9 ortholog. Sci. Adv., 4(10), eaau0766.

Chyou, T.-Y., & Brown, C. M. (2019). Prediction and diversity of tracrRNAs from type II CRISPR-Cas systems. RNA Biol., 16(4), 423–434.

Common, J., Morley, D., Westra, E. R., & van Houte, S. (2019). CRISPR-Cas immunity leads to a coevolutionary arms race between streptococcus thermophilus and lytic phage. Philos. Trans. R. Soc. Lond. B Biol. Sci., 374(1772), 20180098.

Crooks, G. E., Hon, G., Chandonia, J.-M., & Brenner, S. E. (2004). WebLogo: A sequence logo generator. Genome Res., 14(6), 1188–1190.

Deltcheva, E., Chylinski, K., Sharma, C. M., Gonzales, K., Chao, Y., Pirzada, Z. A., Eckert, M. R., Vogel, J., & Charpentier, E. (2011). CRISPR RNA maturation by trans-encoded small RNA and host factor RNase III. Nature, 471(7340), 602–607.

Demirci, S., Leonard, A., Essawi, K., & Tisdale, J. F. (2021). CRISPR-Cas9 to induce fetal hemoglobin for the treatment of sickle cell disease. Mol. Ther. Methods Clin. Dev., 23, 276–285.

Doench, J. G. (2018). Am I ready for CRISPR? a user’s guide to genetic screens. Nat. Rev. Genet., 19(2), 67–80.

Esvelt, K. M., Mali, P., Braff, J. L., Moosburner, M., Yaung, S. J., & Church, G. M. (2013). Orthogonal cas9 proteins for RNA-guided gene regulation and editing. Nat. Methods, 10(11), 1116–1121.

Frangoul, H., Altshuler, D., Cappellini, M. D., Chen, Y.-S., Domm, J., Eustace, B. K., Foell, J., de la Fuente, J., Grupp, S., Handgretinger, R., Ho, T. W., Kattamis, A., Kernytsky, A., Lekstrom-Himes, J., Li, A. M., Locatelli, F., Mapara, M. Y., de Montalembert, M., Rondelli, D., … Corbacioglu, S. (2021). CRISPR-Cas9 gene editing for sickle cell disease and *β*-thalassemia. N. Engl. J. Med., 384(3), 252–260.

Garneau, J. E., Dupuis, M.-È., Villion, M., Romero, D. A., Barrangou, R., Boyaval, P., Fremaux, C., Horvath, P., Magadán, A. H., & Moineau, S. (2010). The CRISPR/Cas bacterial immune system cleaves bacteriophage and plasmid DNA. Nature, 468(7320), 67–71.

Gleditzsch, D., Pausch, P., Müller-Esparza, H., Özcan, A., Guo, X., Bange, G., & Randau, L. (2019). PAM identification by CRISPR-Cas effector complexes: Diversified mechanisms and structures. RNA Biol., 16(4), 504–517.

Harrington, L. B., Burstein, D., Chen, J. S., Paez-Espino, D., Ma, E., Witte, I. P., Cofsky, J. C., Kyrpides, N. C., Banfield, J. F., & Doudna, J. A. (2018). Programmed DNA destruction by miniature CRISPR-Cas14 enzymes. Science, 362(6416), 839–842.

Heler, R., Samai, P., Modell, J. W., Weiner, C., Goldberg, G. W., Bikard, D., & Marraffini, L. A. (2015). Cas9 specifies functional viral targets during CRISPR-Cas adaptation. Nature, 519(7542), 199–202.

Hidalgo-Cantabrana, C., Goh, Y. J., Pan, M., Sanozky-Dawes, R., & Barrangou, R. (2019). Genome editing using the endogenous type I CRISPR-Cas system in lactobacillus crispatus. Proc. Natl. Acad. Sci. U. S. A., 116(32), 15774–15783.

Jinek, M., Chylinski, K., Fonfara, I., Hauer, M., Doudna, J. A., & Charpentier, E. (2012). A programmable dual-RNA-guided DNA endonuclease in adaptive bacterial immunity. Science, 337(6096), 816–821.

Jumper, J., Evans, R., Pritzel, A., Green, T., Figurnov, M., Ronneberger, O., Tunyasuvunakool, K., Bates, R., Žídek, A., Potapenko, A., Bridgland, A., Meyer, C., Kohl, S. A. A., Ballard, A. J., Cowie, A., Romera-Paredes, B., Nikolov, S., Jain, R., Adler, J., … Hassabis, D. (2021). Highly accurate protein structure prediction with AlphaFold. Nature, 596(7873), 583–589.

Kang, Y., Chu, C., Wang, F., & Niu, Y. (2019). CRISPR/Cas9-mediated genome editing in nonhuman primates. Dis. Model. Mech., 12(10), dmm039982.

Karah, N., Samuelsen, Ø., Zarrilli, R., Sahl, J. W., Wai, S. N., & Uhlin, B. E. (2015). CRISPR-cas subtype I-Fb in acinetobacter baumannii: Evolution and utilization for strain subtyping. PLoS One, 10(2), e0118205.

Kim, E., Koo, T., Park, S. W., Kim, D., Kim, K., Cho, H.-Y., Song, D. W., Lee, K. J., Jung, M. H., Kim, S., Kim, J. H., Kim, J. H., & Kim, J.-S. (2017). In vivo genome editing with a small cas9 orthologue derived from campylobacter jejuni. Nat. Commun., 8(1), 14500.

Komor, A. C., Badran, A. H., & Liu, D. R. (2017). CRISPR-based technologies for the manipulation of eukaryotic genomes. Cell, 168(1-2), 20–36.

Leenay, R. T., Maksimchuk, K. R., Slotkowski, R. A., Agrawal, R. N., Gomaa, A. A., Briner, A. E., Barrangou, R., & Beisel, C. L. (2016). Identifying and visualizing functional PAM diversity across CRISPR-Cas systems. Mol. Cell, 62(1), 137–147.

Li, Z., Wang, X., Xu, D., Zhang, D., Wang, D., Dai, X., Wang, Q., Li, Z., Gu, Y., Ouyang, W., Zhao, S., Huang, B., Gong, J., Zhao, J., Chen, A., Shen, Y., Dong, Y., Zhang, W., Xu, X., … Jiang, Y. (2020). DNB-based on-chip motif finding: A high-throughput method to profile different types of protein-DNA interactions. Sci. Adv., 6(31), eabb3350.

Makarova, K. S., Wolf, Y. I., Iranzo, J., Shmakov, S. A., Alkhnbashi, O. S., Brouns, S. J. J., Charpentier, E., Cheng, D., Haft, D. H., Horvath, P., Moineau, S., Mojica, F. J. M., Scott, D., Shah, S. A., Siksnys, V., Terns, M. P., Venclovas, Č., White, M. F., Yakunin, A. F., … Koonin, E. V. (2020). Evolutionary classification of CRISPR-Cas systems: A burst of class 2 and derived variants. Nat. Rev. Microbiol., 18(2), 67–83.

Mao, Y., Botella, J. R., Liu, Y., & Zhu, J.-K. (2019). Gene editing in plants: Progress and challenges. Natl. Sci. Rev., 6(3), 421–437.

Mendoza, B. J., & Trinh, C. T. (2018). In silico processing of the complete CRISPR-Cas spacer space for identification of PAM sequences. Biotechnol. J., 13(9), e1700595.

Mitrofanov, A., Ziemann, M., Alkhnbashi, O. S., Hess, W. R., & Backofen, R. (2022). CRISPRtracrRNA: Robust approach for CRISPR tracrRNA detection. Bioinformatics, 38(Suppl_2), ii42–ii48.

Mu, Y., Zhang, C., Li, T., Jin, F.-J., Sung, Y.-J., Oh, H.-M., Lee, H.-G., & Jin, L. (2022). Development and applications of CRISPR/Cas9-based genome editing in lactobacillus. Int. J. Mol. Sci., 23(21), 12852.

Musunuru, K., Chadwick, A. C., Mizoguchi, T., Garcia, S. P., DeNizio, J. E., Reiss, C. W., Wang, K., Iyer, S., Dutta, C., Clendaniel, V., Amaonye, M., Beach, A., Berth, K., Biswas, S., Braun, M. C., Chen, H.-M., Colace, T. V., Ganey, J. D., Gangopadhyay, S. A., … Kathiresan, S. (2021). In vivo CRISPR base editing of PCSK9 durably lowers cholesterol in primates. Nature, 593(7859), 429–434.

Pausch, P., Al-Shayeb, B., Bisom-Rapp, E., Tsuchida, C. A., Li, Z., Cress, B. F., Knott, G. J., Jacobsen, S. E., Banfield, J. F., & Doudna, J. A. (2020). CRISPR-CasΦ from huge phages is a hypercompact genome editor. Science, 369(6501), 333–337.

Pickar-Oliver, A., & Gersbach, C. A. (2019). The next generation of CRISPR-Cas technologies and applications. Nat. Rev. Mol. Cell Biol., 20(8), 490–507.

Pyne, M. E., Bruder, M. R., Moo-Young, M., Chung, D. A., & Chou, C. P. (2016). Harnessing heterologous and endogenous CRISPR-Cas machineries for efficient markerless genome editing in clostridium. Sci. Rep., 6, 25666.

Qin, Z., Yang, Y., Yu, S., Liu, L., Chen, Y., Chen, J., & Zhou, J. (2021). Repurposing the endogenous type I-E CRISPR/Cas system for gene repression in gluconobacter oxydans WSH-003. ACS Synth. Biol., 10(1), 84–93.

Ran, F. A., Cong, L., Yan, W. X., Scott, D. A., Gootenberg, J. S., Kriz, A. J., Zetsche, B., Shalem, O., Wu, X., Makarova, K. S., Koonin, E. V., Sharp, P. A., & Zhang, F. (2015). In vivo genome editing using staphylococcus aureus cas9. Nature, 520(7546), 186–191.

Russel, J., Pinilla-Redondo, R., Mayo-Muñoz, D., Shah, S. A., & Sørensen, S. J. (2020). CRISPRCasTyper: Automated identification, annotation, and classification of CRIS Cas loci. CRISPR J., 3(6), 462–469.

Rybnicky, G. A., Fackler, N. A., Karim, A. S., Köpke, M., & Jewett, M. C. (2022). Spacer2PAM: A computational framework to guide experimental determination of functional CRISPR-Cas system PAM sequences. Nucleic Acids Res., 50(6), 3523– 3534.

Sayers, E. W., Beck, J., Bolton, E. E., Bourexis, D., Brister, J. R., Canese, K., Comeau, D. C., Funk, K., Kim, S., Klimke, W., Marchler-Bauer, A., Landrum, M., Lathrop, S., Lu, Z., Madden, T. L., O’Leary, N., Phan, L., Rangwala, S. H., Schneider, V. A., … Sherry, S. T. (2021). Database resources of the national center for biotechnology information. Nucleic Acids Res., 49(D1), D10–D17.

Sun, A., Li, C.-P., Chen, Z., Zhang, S., Li, D.-Y., Yang, Y., Li, L.-Q., Zhao, Y., Wang, K., Li, Z., Liu, J., Liu, S., Wang, J., & Liu, J.-J. G. (2023). The compact Cas*π* (cas12l) ‘bracelet’ provides a unique structural platform for DNA manipulation. Cell Res., 33(3), 229–244.

Tang, L. C., & Gu, F. (2020). Next-generation CRISPR-Cas for genome editing: Focusing on the cas protein and PAM. Yi Chuan, 42(3), 236–249.

Vale, P. F., & Little, T. J. (2010). CRISPR-mediated phage resistance and the ghost of coevolution past. Proc. Biol. Sci., 277(1691), 2097–2103.

Walker, J. E., Lanahan, A. A., Zheng, T., Toruno, C., Lynd, L. R., Cameron, J. C., Olson, D. G., & Eckert, C. A. (2020). Development of both type I-B and type II CRISPR/Cas genome editing systems in the cellulolytic bacterium clostridium thermocellum. Metab. Eng. Commun., 10(e00116), e00116.

Wang, J., Li, J., Zhao, H., Sheng, G., Wang, M., Yin, M., & Wang, Y. (2015). Structural and mechanistic basis of PAM-dependent spacer acquisition in CRISPR-Cas systems. Cell, 163(4), 840–853.

Wang, J. Y., & Doudna, J. A. (2023). CRISPR technology: A decade of genome editing is only the beginning. Science, 379(6629), eadd8643.

Xiao, Y., Ng, S., Nam, K. H., & Ke, A. (2017). How type II CRISPR-Cas establish immunity through Cas1-Cas2-mediated spacer integration. Nature, 550(7674), 137– 141.

Xu, C., Zhou, Y., Xiao, Q., He, B., Geng, G., Wang, Z., Cao, B., Dong, X., Bai, W., Wang, Y., Wang, X., Zhou, D., Yuan, T., Huo, X., Lai, J., & Yang, H. (2021). Programmable RNA editing with compact CRISPR-Cas13 systems from uncultivated microbes. Nat. Methods, 18(5), 499–506.

Xue, C., & Greene, E. C. (2021). DNA repair pathway choices in CRISPR-Cas9-mediated genome editing. Trends Genet., 37(7), 639–656.

Zetsche, B., Gootenberg, J. S., Abudayyeh, O. O., Slaymaker, I. M., Makarova, K. S., Essletzbichler, P., Volz, S. E., Joung, J., van der Oost, J., Regev, A., Koonin, E. V., & Zhang, F. (2015). Cpf1 is a single RNA-guided endonuclease of a class 2 CRISPR-Cas system. Cell, 163(3), 759–771.

Zhang, J., Zong, W., Hong, W., Zhang, Z.-T., & Wang, Y. (2018). Exploiting endogenous CRISPR-Cas system for multiplex genome editing in clostridium tyrobutyricum and engineer the strain for high-level butanol production. Metab. Eng., 47, 49–59.

Zuker, M. (2003). Mfold web server for nucleic acid folding and hybridization prediction. Nucleic Acids Res., 31(13), 3406–3415.

